# VgeneRepertoire.org identifies and stores variable genes of immunoglobulins and T-cell receptors from the genomes of jawed vertebrates

**DOI:** 10.1101/002139

**Authors:** David N. Olivieri, Francisco Gambón-Deza

## Abstract

The VgeneRepertoire.org platform (http://vgenerepertoire.org) is a new public database repository for variable (V) gene sequences that encode immunoglobulin and T-cell receptor molecules. It identifies the nucleic and amino acid sequences of more than 20,000 genes, providing their exon location in either the contig, scaffold, or chromosome region, as well as locus information for more than 100 jawed vertebrate taxa whose genomes have been sequenced. This web repository provides support to immunologists interested in these molecules and aids in comparative phylogenetic studies.

## 1. Introduction

The increasing availability of sequenced genomes from a wide array of jawed vertebrate taxa requires intensive data analysis for a comprehensive understanding of constituent genes, their function, and their evolution (Flajnik, 2002; Rothenfluh et al., 1995). In Comparative Immunology, efforts to use this WSG data to study the genes that code for immunoglobulins (Ig) and T-cell receptors (TCR), are still in their infancy. Of particular interest are those genes that encode parts of Ig and TCR molecules responsible for antigen recognition. The basic genomic elements of the adaptive immune system (AIS) make it possible to produce an enormous variability with high specificity, through a recombination process using different domain types, referred to as the V (variable), D (differentiation), and J (joining) regions (Hozumi & Tonegawa, 1976). Of these domain types, the largest are the V-regions sequences. Thus, one of the principal factors governing the specie's capacity to recognize antigen, is the size and structure of the V-gene repertoire within its genome. Comparative studies of the V-gene repertoire between taxa is essential for a obtaining a comprehensive understanding of the mechanisms by which the AIS communicates with the environment]. In particular, clues about gene duplication and birth/death processes can be obtained by contrasting the evolution of these V-gene sequences in different extant vertebrates which have been influenced by separate ecological niches.

VgeneRepertoire.org is a Web platform and public repository consisting of V-gene sequences identified from the genomes of jawed vertebrates that have been made available at the NCBI. These V-gene sequences were obtained from Whole Shotgun Genome (WSG) data sets with the *VgenExtractor* bioinformatics tool, which has a recognition accuracy of approximately 95%. Until now, complete V-gene annotations have only been available in two species (i.e. human and mouse), while incomplete annotations have existed in a few species, and even in these cases, only within certain loci. The *VgeneRepertoire.org* repository represents a comprehensive collection of V-genes (approximately 20,000 to date) from a wide array of taxa (approximately 100; from the orders mammals, reptiles, birds, and fish).

For each taxon, sequences derived from several different WSG datasets or assembly versions are possible (and indeed the case for some species). For a given WSG of a species, information pertaining to each Ig or TCR V-gene sequence includes: (a) the contig region (given by the GID and a hyperlink to the GeneBank repository through its accession number), (b) the nucleic and amino acid sequences, and (c) the locus isotype (i.e., IGHV, IGLV, IGKV, TRA/DV, TRBV, etc) (Tonegawa et al., 1 Jan. 1978; Davis & Bjorkman, 1988). Also for a particular WSG dataset of an organism, information is provided about the sequencing experiment, the BioProjects code, and a summary of the V-genes sequence distribution amongst the different loci. For selecting V-gene sequences, many search options are available, including: (1) searches by single/multiple loci by species, order, or family, (2) searches by single/multiple species depending upon order or subfamily, or (3) selecting by browsing through the entire collection of taxa. In all cases, single or multiple sequences may be selected and exported to Fasta format, or exported to one of the tools included in the Web platform for phylogenetic analysis.

The workflow management pipeline of the *VgeneRepertoire.org* consist of several custom bioinformatic tools developed by our group for maintaining and curating the database. The first stage of the pipeline is a curation stage that determines whether new WSG datasets are available, and if so, produce the commands necessary for obtaining the corresponding sequence files. The next stage extracts V-gene sequences from the WSG using *VgeneExtractor* Olivieri et al. (2013). For each V-gene encountered, we predict the most probable locus to which it belongs by using a new machine learning tool, *VgeneClassify*. Finally, we perform phylogenetic analysis using Clustal-omega (Sievers F, 2014) and Fasttree (Price et al., 2009, 2010) to validate the classification of the previous step and to confirm cladistic relations between sequences of each loci. The pipeline for obtaining sequences as well as a basic workflow of the web platform for searching V-gene sequences is shown in Figure 1.

**Figure 1.**
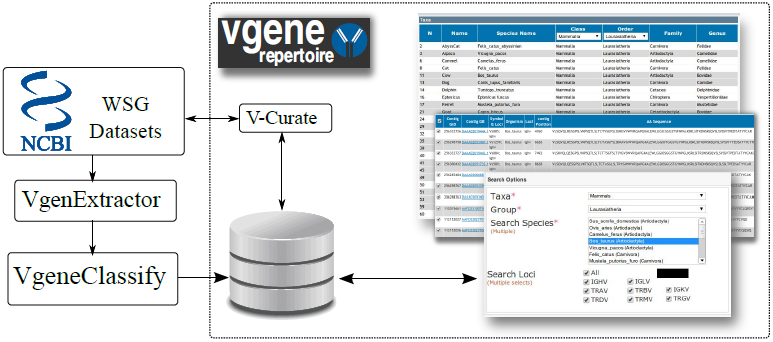
System architecture.

A detailed discussion motivating the study of V-genes across a wide spectrum of taxa can be found elsewhere Gambón-Deza et al. (2012); Olivieri et al. (2013). Briefly, comparative evolutionary studies of these V-genes can also unveil broader clues about habitat adaptation from the different antigen recognition mechanisms of Ig as compared to TCR. In this respect, both Ig and TCR, although possessing similar V-gene structures, behave and function quite differently. While, antibody (Ig molecules) are able to recognize antigen molecules directly in soluble form (when secreted massively by of B-cell populations possessing these specific Ig receptors), the TCR molecules are restricted by a complex interplay with the major histocompatibility complex (MHC) and the antigenic protein to be recognized (Huppa & Davis, 2003). Such differences have profound implications that govern the origin and evolution of the genes that encode these immuno-logic molecules.

Here we describe some of the details of the pipeline and workflow management software used to curate the *VgeneRepertoire.org* repository. We provide a summary of the V-genes available in the repository. This data contains a plethora of information and should motivate future studies that will unravel important clues about the evolution of the adaptive immune system through multispecies studies of gene duplication processes as well as birth/death rates between these loci.

## 2. Methods

In both antibodies as well as in TCR, the antigen recognition is always mediated by its variable regions, which are composed of two polypeptide chains encoded by the V-genes. The V-gene exon sequences are found with the *VgenExtractor* software tool Olivieri et al. (2013) by marching through the genome sequence file and identifying specific nucleic acid patterns, called *recombination signal sequence* (RSS), that mark the start location of the V region (i.e., located at the downstream 3′ end of the V-exon boundary). These RSS markers are composed of the same nucleic acid sequence in all loci (with a consensus of the form, CACAGTG), and are known to be universal for all vertebrates, making their identification consistent across isotypes and species. For each occurrence of sequences identical to an RSS, the upstream exon region is tested to determine whether it could be a candidate V-gene sequence using the following criteria: it has an approximate length of 300 bp, the reading frame is unaltered, and in the translation to amino acids, it contains two canonical cysteines and a tryptophan at conserved positions within the exon. Psuedogenes, which are exons containing alterations in the reading frame or stop codons during the translation, are not considered further in the analysis and are discarded.

Apart from identifying functional V-genes, it is important to identify their isotype, or equivalently the chain of the Ig or TCR molecules which they encode. Because WSG datasets do not have assembled chromosome, the locus is unknown and must be inferred from sequence comparisons or molecular phylogenetic (i.e., cladistic) relationships. In the adaptive immune system of jawed vertebrates, it is known that there exist at least seven separate loci in the genome corresponding to Ig and TCR chains. In mammals, three V-gene loci correspond to immunoglobulins: one heavy chain (IGHV), and two light chains, referred to as *κ* (IGKV) and *λ* (IGLV). The other loci in mammals correspond to TCR chains: the *α/β* and the *γ/δ*. The TCR *α/β* is composed of two chains (*α* y *β*), in which the variable region of each chain is coded by a different loci (TRAV and TRBV, respectively). The TCR *γ/δ* also consists of two chains (the *γ* and *δ*), where their variable regions are encoded by two different loci (TRGV and TRDV). The loci TRDV has the same chromosomal location as that of the TRAV.

For a given V-gene sequence, our *VgeneClassify* software tool determines the isotype, or locus, by using a probabilistic model, which uses feature vectors formed from the output of a Blastp sequence similarity search against the entire protein database. The feature vector extracts information about the set of identity scores, sequence coverages, and text descriptions. A likelihood model uses these feature vectors to assign the candidate sequence to one of the V-gene loci types. The resulting loci classification has been validated and is consistent with a phylogenetic analysis of all the sequences of a species, where each locus forms a separate clade in the tree.

A summary of the method is shown in the block diagram of Figure 2 for two example sequences. In the figure, The example sequence 1 (upper, yellow) is processed with Blastp and the output (upper, in blue) shows a listing with low identity/coverage as well as protein descriptions that contain many stop words. The algorithm processes this information and forms a feature vector, that is used in a model to generate a likelihood estimate for pertaining to any one of the V-gene loci. This sequence will produce a low likelihood for all loci classes and be discarded. On the other hand, the example sequence 2 (lower, yellow) produces a blastp output that contains high content words, as well as high identity/coverage scores. The feature vector for this sequence will produce a high likelihood for the Ig heavy chain locus. The result of the loci prediction is validated from a phylogenetic analysis of the V-gene sequences, where each locus occupies a separate clade (as seen in the example of Figure 2).

**Figure 2.**
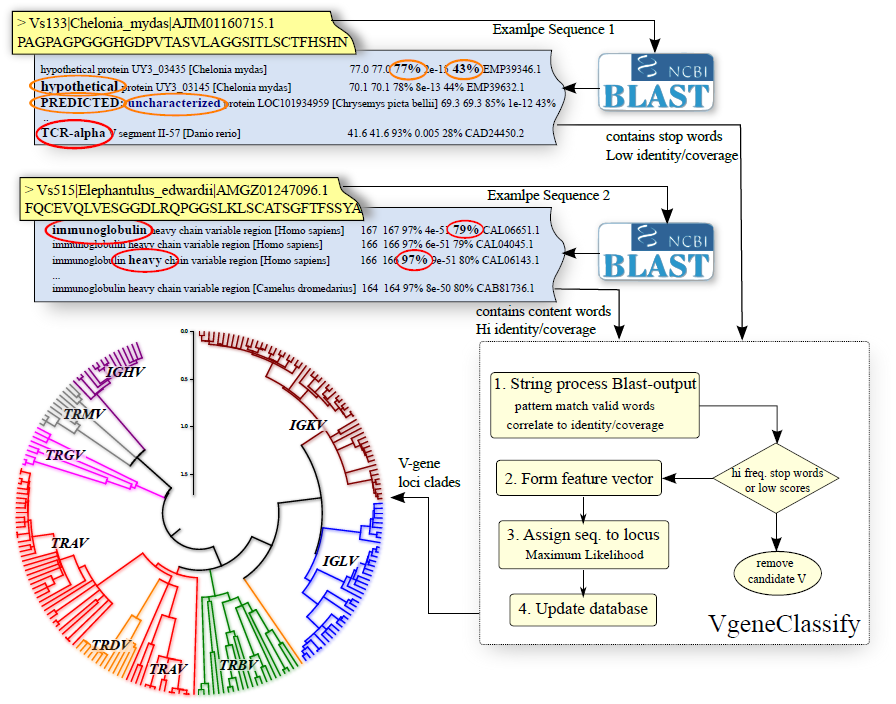
Block diagram of machine learning loci determination. Two example sequences are used to illustrate the operation of the algorithm. A feature vector is constructed from the Blastp output of a sequence similarity search against the protein database. Likelihoods are assigned to each possible V-gene locus. A sequence that produces stop words and low identity/coverage scores will have low likelihood for all loci and be discarded.

The different modules in the workflow pipeline are executed through a set of custom python classes. The design is modular and computationally efficient, allowing for the addition of other analysis methods in the future. The results of V-gene discovery and classification pipeline from WSG datasets are automatically loaded into the PostgreSQL database. The Web server is written in PHP and uses AJAX/JQuery for displaying and manipulating data loaded into the Web browser.

## 3. Results

We have indexed over 20000 V-genes from WSG datasets of mammals, reptiles, birds, and fish. Figure 3 shows a summary of mean V-gene number within each order/family of mammals and reptiles included in the database to date. In this summary, 19643 V-genes from 82 mammal and 11 reptile species are represented, which are the number of WSG presently available to date, at the time of writing this article.

**Figure 3.**
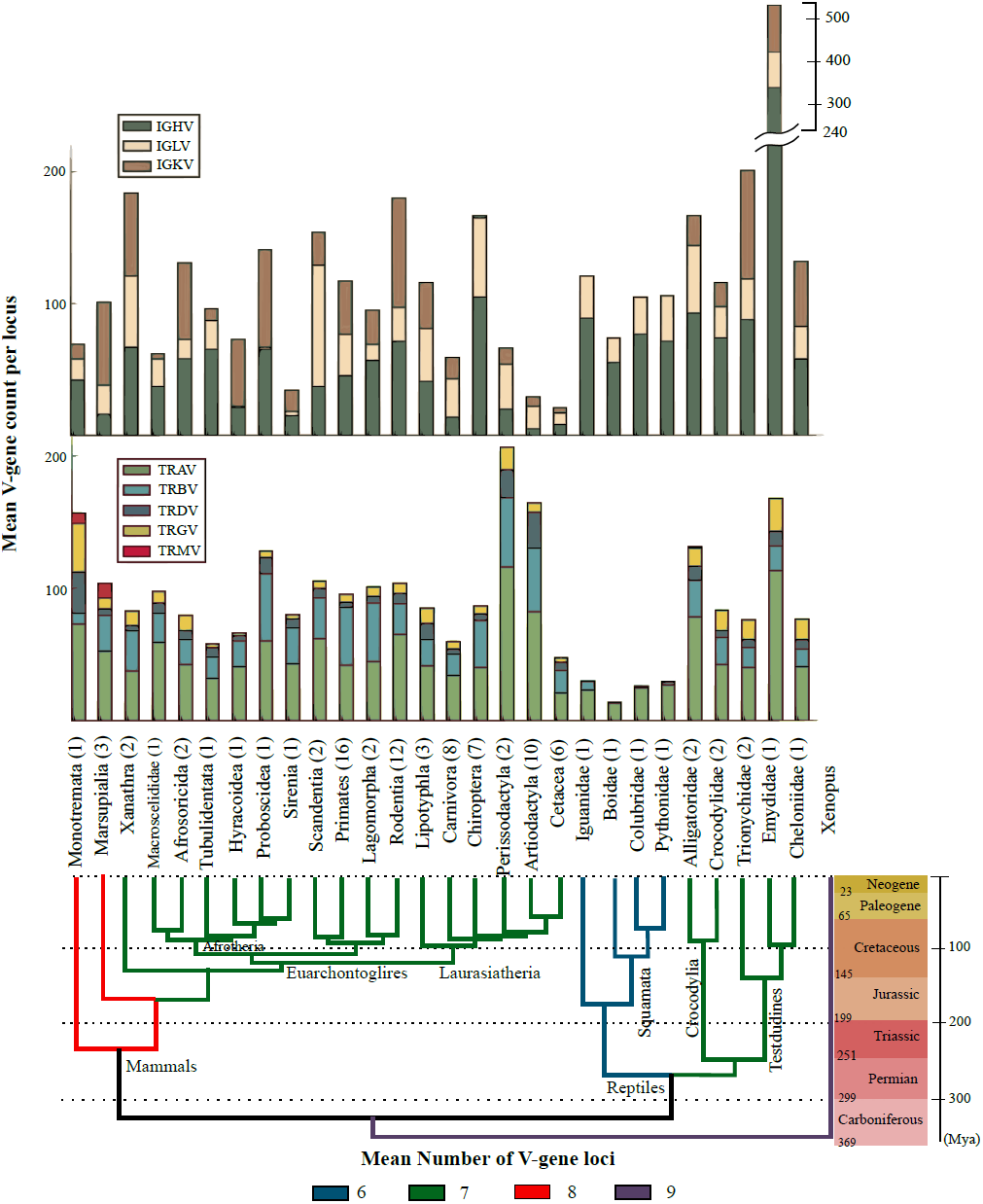
Repertoire of Taxa. The time scales of speciation events in the phylogenetic tree were taken from (O'Leary et al., 2013; Hedges & Kumar, 2009).

From Figure 3, quantitative information about V-genes, across a many orders/families, can be obtained. The figure plots the Ig loci on the upper barplot and the TCR loci in the lower barplot. In nearly all families/orders, the Ig heavy chain and the TCR*α* chains predominate over other loci. As seen, the mean number of V-genes in the Testudine family, Emydidae, is larger than found in other species (shown off-scale in the figure). Also, it is seen that Squamata possess few TCR V-genes but approximately 100 IG V-genes (which is the average across all species) but lack the Ig *κ* chain. To the contrary, the Perissodactyla and Artiodactyla have a relatively large mean number of TCR V-genes but a smaller amount of Ig V-genes. A similar trend between Ig and TCR can be seen in other families/ordens, to a lesser degree, except in the Cetacea (comprised of six species in this study), which have few V-genes in both the Ig and TCR loci.

We have plotted the above data together with the phylogenetic relations between the different families/orders since it can reveal more information. The phylogenetic tree was constructed using speciation event data taken from (O'Leary et al., 2013; Hedges & Kumar, 2009). We colored the branches of the tree depending upon the mean number of loci within each family. As can be seen, most mammals have seven loci in total, except for the Monotremata and Marsupialia, which possess 8 loci (possessing the extra TRM locus). Our previous studies have indicated that the Squamata have lost the V*κ* chain, possibly due to genomic alterations in its more distant relative, the Anole. Finally, Xenopus has the most mean loci, since it also possess the Ig*σ*.

Figure 4 show the cummulative number of V-gene sequences in the database (filled blue background; right y-axis) as well as the total number sequences for each family/order for mammals and reptiles.

**Figure 4.**
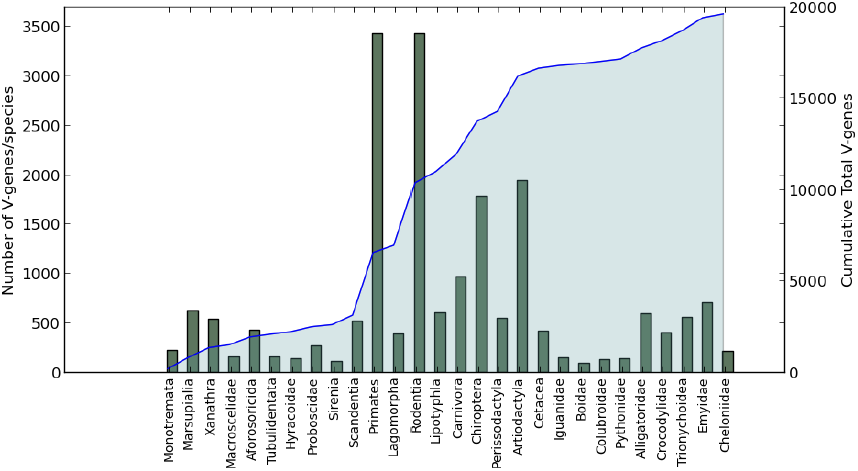
Total cumulative number of V-genes in the database.

## 4. Discussion

The variability of the Ig and TCR V-gene repertoire amongst different species is still not understood (Flajnik & Kasahara, 2010; Ota & Nei, 1994; Das et al., 2012). While most species possess a total number of V-genes between 50 and 300, some species possess more than double this amount (e.g. microbat, rat, and Chrysemys picta belli). While each jawed vertebrate species has an adaptive immune system derived from the same origins and functioning under similar mechanisms, the amount of basic genetic material available for recognizing antigen through recombination processes appears to be different. Despite these differences, there are some species that can apparently make a sufficiently effective immune response with very few V-genes, while other species have a massive amount of V-genes, presumably through some genetic events in their evolution that have favored a large amount of such genes.

The VgeneRepertoire.org Web platform and repository can begin to address such questions. Structural studies of the protein products from these V-genes may also provide clues hidden in their sequences. These sequences consist of different regions: some provide the fundamental physical structure (called framework regions), while others are directly related to antigen recognition (referred to as complementarity-determining regions, CDRs). For the first time, this database provides the opportunity to study these regions with data from more than 80 mammals and 11 reptiles, mining potential biological relationships hereto unknown.

With the data from VgeneRepertoire.org, we have witnessed trends with respect to the total number of V-genes amongst a wide array of taxa. Two phenomena may be at work. If a V-region loses the capacity to recognize a particular antigen through a point mutation in the sequence, there may be an associated gain for recognition of another antigen. Although increasing the number of V-genes may have been an evolutionary strategy increase the effectiveness of an immune response, it is known just through an increase in V-genes is not a sufficient condition. Indeed, additional mechanism are operating, such as somatic mutation, that must be affected to in order to increase the diversity for recognizing antigen (Tomlinson et al., 1996).

The specific diversification processes of these genes and subsequent changes between species can help in the understanding of evolutionary processes that guided these species. These genes are converted into structures that modify the language of communication with the outside world of molecular structures. The sequences of this database are able to decipher this language.

